# A biologically inspired repair mechanism for neuronal reconstructions with a focus on human dendrites

**DOI:** 10.1101/2023.06.15.545190

**Authors:** Moritz Groden, Hannah M. Moessinger, Barbara Schaffran, Javier DeFelipe, Ruth Benavides-Piccione, Hermann Cuntz, Peter Jedlicka

## Abstract

Investigating and modelling the functionality of human neurons remains challenging due to the technical limitations, resulting in scarce and incomplete 3D anatomical reconstructions. Here we used a morphological modelling approach based on optimal wiring to repair the parts of a dendritic morphology that were lost due to incomplete tissue samples. In *Drosophila*, where dendritic regrowth has been studied experimentally using laser ablation, we found that modelling the regrowth reproduced a bimodal distribution between regeneration of cut branches and invasion by neighbouring branches. Interestingly, our repair model followed growth rules similar to those for the generation of a new dendritic tree. To generalise the repair algorithm from *Drosophila* to mammalian neurons, we artificially sectioned reconstructed dendrites from mouse and human hippocampal pyramidal cell morphologies, and showed that the regrown dendrites were morphologically similar to the original ones. Furthermore, we were able to restore their electrophysiological functionality, as evidenced by the recovery of their firing behaviour. Importantly, we show that such repairs also apply to other neuron types including hippocampal granule cells and cerebellar Purkinje cells. We then extrapolated the repair to incomplete human CA1 pyramidal neurons, where the anatomical boundaries of the particular brain areas innervated by the neurons in question were known. Interestingly, the repair of incomplete human dendrites helped to simulate the recently observed increased synaptic thresholds for dendritic NMDA spikes in human versus mouse dendrites. To make the repair tool available to the neuroscience community, we have developed an intuitive and simple graphical user interface (GUI), which is available in the *TREES Toolbox* (www.treestoolbox.org).

**In brief:** We use morphological modelling inspired by the regeneration of various artificially cut neuron types and repair incomplete human and nonhuman neuronal dendritic reconstructions.

**Author summary:** Reconstructing neuronal dendrites by drawing their 3D branching structures in the computer has proven to be crucial for interpreting the flow of electrical signals and therefore the computations that dendrites implement on their inputs. These reconstructions are tedious and prone to disruptive limitations imposed by experimental procedures. In recent years, complementary computational procedures have emerged that reproduce the fine details of morphology in theoretical models. These models allow, for example, to populate large-scale neural networks and to study structure-function relationships. In this work we use a morphological model based on optimised wiring for signal conduction and material cost to repair faulty reconstructions, in particular those of human hippocampal dendrites, which are rare and precious but often cut due to technical limitations. Interestingly, we find that our synthetic repair mechanism reproduces the two distinct modes of repair observed in real dendrites: regeneration from the severed branch and invasion from neighbouring branches. Our model therefore provides both a useful tool for single-cell electrophysiological simulations and a useful theoretical concept for studying the biology of dendrite repair.

**Highlights:**

- Optimal wiring-based growth algorithm replicates regrowth of artificially cut dendrites
- The growth algorithm repairs cut dendrites in incomplete reconstructions
- The algorithm works for diverse neuron types in multiple species
- The repair of morphology restores original electrophysiology
- The repair of morphology supports simulations of high synaptic thresholds for NMDA spikes in human dendrites
- The repair tool with user interface is available in the *TREES Toolbox*

## Introduction

It is well established that dendritic geometry affects neuronal function (London and Häusser, 2005; Stuart *et al*., 2016; Poirazi and Papoutsi, 2020; Platschek *et al*., 2016; Zhu *et al*., 2016). For example, a change in dendritic size or topology may significantly alter the neuronal firing behaviour (Mainen and Sejnowski, 1996; Bekkers and Häusser, 2007; van Ooyen *et al*., 2002; van Elburg and van Ooyen, 2010) in a possibly selective manner (Park *et al*., 2019). Several studies on the morphology and electrophysiology of human neurons have revealed their specific enhanced computational features (Beaulieu-Laroche *et al*., 2018; Gidon *et al*., 2020; Testa-Silva *et al*., 2022; Elston *et al*., 2001). However, systematic investigations of the relationship between the structure and the electrophysiological properties of human dendrites in computational models remain challenging (Fisek and Häusser, 2020; Segev and London, 2000) since complete 3D reconstructions are scarce (DeFelipe, 2015). The sparse anatomical data that is available usually comes from both autopsies of healthy donors and biopsies of patients with brain diseases such as epilepsy or brain tumours (Buchin *et al*., 2020; Domínguez-Álvaro *et al*., 2018; Palacios Bote *et al*., 2008). These diseases can significantly alter the morphology and electrophysiology of a neuron (Houser, 1992; Glass and Dragunow, 1995), resulting in severely impaired cognitive function (Shuman *et al*., 2017). Such pathological dendritic data may limit scientific conclusions if they are interpreted as coming from healthy controls.

Moreover, due to the large size of human neurons and the technical limitations, the reconstruction process is susceptible to errors, often leaving the morphology reconstructions incomplete (Hamam and Kennedy, 2003; Glaser and Van der Loos, 1981; Benavides-Piccione *et al*., 2020, see for example in **Figure 1A, B** and **C**). In addition, staining dyes injected into larger neurons often fail to reach the most distal dendritic areas (De Schutter and Jaeger, 2000; Horcholle-Bossavit *et al*., 2000). Such incomplete reconstructions, further limit the ability to study dendritic anatomy. However, the characterisation of morphological differences between human and other species’ neural circuits (Benavides-Piccione *et al*., 2020; Mihaljevic *et al*., 2020; Mihaljević *et al*., 2021) is of great importance, since they have been shown to lead to distinct computational properties (Kö tter and Feizelmeier, 1998; Eyal *et al*., 2016; Beaulieu-Laroche *et al*., 2021; Elston *et al*., 2001). Therefore, more complete human morphologies are urgently needed for a better understanding of human neuronal physiology and pathophysiology and for the creation of realistic computational models of human dendrites (Hunt *et al*., 2022; Gidon *et al*., 2020; Segev and London, 2000; Poirazi and Papoutsi, 2020).

**Figure 1.**
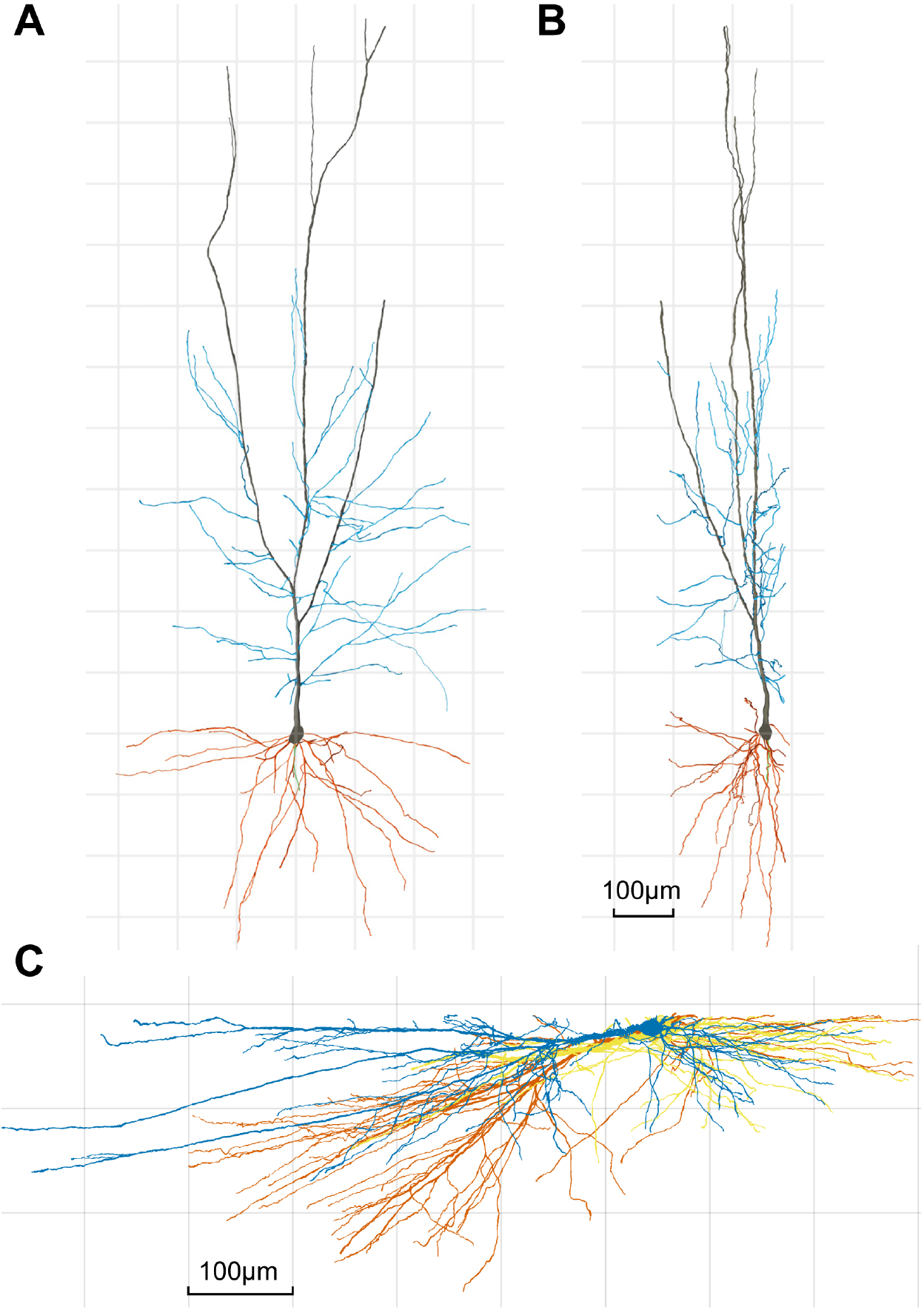
Examples of human CA1 pyramidal reconstructions that were cut in the same plane during tissue sectioning. Example of a 3D-reconstructed human CA1 pyramidal cell shown on the XY **A**, and YZ **B**, planes, to illustrate that, due to technical limitations, part of the dendritic arbor closest to the surface of the slice from which the cell soma is injected (typically at a depth of *∼* 30*µm* from the surface) is lost. Axon, main apical, collateral and basal dendrites are shown in green, black, blue and orange, respectively. Scale bar (in panel **B**) = 100*µm*. **C**, Three human CA1 pyramidal neuron reconstructions (yellow, orange and blue) from the same preparation viewed from the side. Raw data from Benavides-Piccione *et al*. (2020).

Solutions to implement repair tools for morphological neuronal reconstructions have been proposed in the past (Abdellah *et al*., 2018). These models usually focused on detecting and fixing or removing artefacts that may occur during the reconstruction process, such as neurites that are not properly connected to the soma, removing segments with zero length, or adjusting dendrites that cross each other (*NeuronR*, Anwar *et al*., 2009). Other morphological growth models are usually implemented as stochastic procedures based on branch probabilities and the number of branching events (Ascoli and Krichmar, 2000; van Pelt and Schierwagen, 2004; Donohue and Ascoli, 2008). These branch probabilities are sampled from experimental distributions. This results in a large number of model parameters that must be adjusted to generate different cell types. Adding entirely new branches to existing dendritic trees is not part of such tools.

For these reasons, in this work we investigated whether *in silico* dendritic growth algorithms based on optimal wiring (Cuntz *et al*., 2010; Baltruschat *et al*., 2020; Ferreira Castro *et al*., 2020) are also able to complete incomplete morphology reconstructions by adding missing parts of the dendritic tree that resemble real structures. Optimal wiring principles allow the dendritic structure to be described by locally optimised graphs, in which total length and path length are minimised (Cuntz *et al*., 2007; Wen and Chklovskii, 2008). An algorithm that weights these two factors by a balancing factor *bf* can generate synthetic trees that closely resemble biological dendrites (Cuntz *et al*., 2010, 2011). Once target points are distributed within a cell-type specific dendritic density field, they can be connected to a tree structure according to these optimised wiring costs in *e.g.* fly (Cuntz *et al*., 2008) or mouse (Cuntz, 2012) as well as in some axons (Budd *et al*., 2010). Given the general applicability of the method, here we investigate whether such morphological modelling can also be used to better understand and implement dendrite repair.

The biological system that inspired our regrowth algorithm was the nervous system of the *Drosophila* larva with so-called da (dendritic arborisation) neurons (Bodmer and Jan, 1987). These are divided into four classes based on their dendritic pattern, classes I–IV. Class IV da neurons grow predominantly in a two-dimensional space (Han *et al*., 2012) and are well known to regrow their dendrites after dendriotomy (Song *et al*., 2012; Stone *et al*., 2014). Almost 98% of all proximally lesioned dendrites showed regrowth, as measured by receptive field coverage after lesioning. Interestingly, in some cases the cut dendrite regenerated from the site of its lesion, and in others the field was covered by invading neighbouring branches of the same neuron, showing a bimodal distribution of dendrite regrowth (Song *et al*., 2012).

In the work presented here, we report that our synthetic growth algorithm has the ability to mimic biological regrowth and to reproduce its two observed modes. In addition, regrowth can be tuned to emerge exclusively from the known incomplete ends of severed dendrite morphologies. Taking advantage of these features, we build a *TREES Toolbox* function *fix tree* and a user interface *fix tree UI* to complete dendritic reconstructions inspired by biological regrowth.

## Results

### A repair mechanism inspired from biology

To develop a repair algorithm for incomplete/damaged dendrites of nerve cells based on biologically inspired mechanisms, we first simulated and analysed the synthetic regrowth of dendrites characterised in a controlled experimental setting. Class IV da neurons of *Drosophila* larvae are a useful and well-studied experimental model system to investigate dendritic growth following dendriotomy (Song *et al*., 2012; Li *et al*., 2018; Stone *et al*., 2014). To simulate the repair mechanism, seven reconstructions of class IV neurons (Ziegler *et al*., 2017) were taken from *NeuroMorpho.org* (Ascoli *et al*., 2007; Parekh and Ascoli, 2013). The location where the cell was cut was chosen as a random branch point of the original morphology. The root of the severed branch (including the branch point) was used as a reference to determine the type of regrowth following the lesion. Branches growing back from this node during the repair process were defined as regenerated. Branches that innervated the lesioned area but did not originate from the lesion node, were considered to be invading the space made available by the lesion.

We implemented a regrowth protocol, using newly distributed target points within the region of the severed branch, and replicated the stochastic regrowth *in silico* (see Methods). Regrowth based solely on optimal wiring principles, balancing path length and total wiring cost (Cuntz *et al*., 2010), successfully reproduced the main features of the dendrites. Importantly, our model replicated the experimentally observed bimodal distribution of branch regeneration versus invasion from neighbouring branches:

Sometimes, the synthetic regrowth invaded the available space with new branches emerging from adjacent branches (**Figure 2A**, compare green branches with severed branches in magenta). At other times, the repair algorithm showed complete regeneration of the lesioned area by a branch originating from the severed node, as shown in the example in **Figure 2B** with similar colours. These two types or modes of synthetic regrowth were in close agreement with similar observations in the experiment of Song *et al*. (2012), indicating that optimal wiring principles may be sufficient to explain these experimental observations (**Figure 2A, B**).

**Figure 2.**
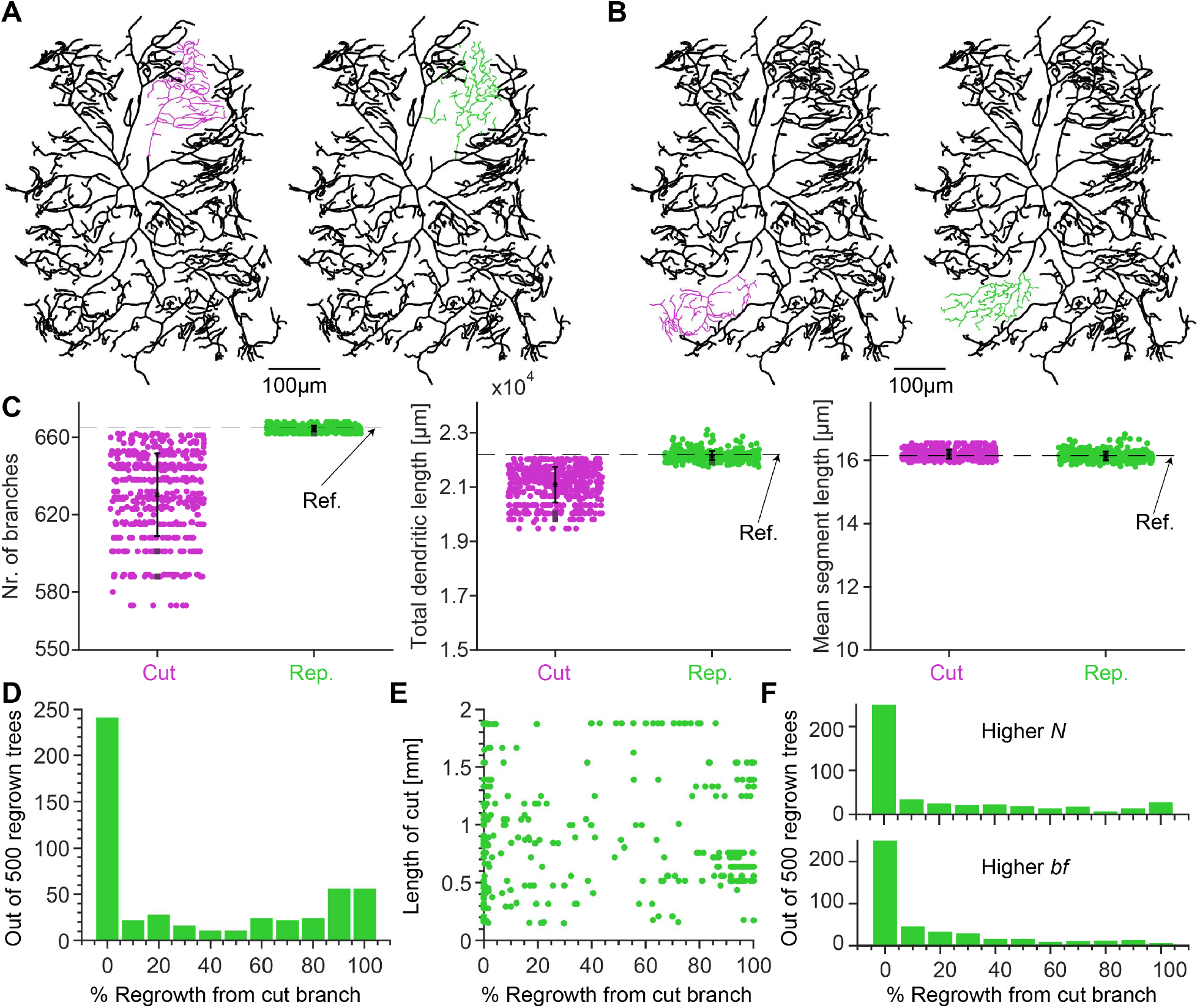
Reproduction of biological regrowth of severed Class IV *Drosophila* neurons. **A** and **B**, *Left*, Reference *Drosophila* larva Class IV morphology with deliberately severed branches marked in magenta. **A**, *Right*, Example of repaired dendrite where invasion has occurred from adjacent branches marked (green). **B**, Sample repair where the severed branch regenerated from the cut end (green). **C**, Morphological statistics of the regrown dendrites from **A** and **B** (green) and 498 other random cuts. The repaired morphologies were compared to the original reference neuron (black + magenta in **A** and **B**) shown here as the black dashed line, as well as to the cut dendrites (magenta). The examples shown in **A** and **B** are represented by the darker square data points. **D**, Histogram for 500 regrown dendrites using our repair function, showing the percentage of regenerated branches. **E**, Percentage of regenerated branches as a function of the size of the removed branch. **F**, Histograms for 500 regrown morphologies as in **D** but for a higher number of target points *N* and a higher balancing factor *bf*.

Reconstructions from *NeuroMorpho.org* allowed us to generate synthetic cells that matched the branching statistics of class IV da neurons (Nanda *et al*., 2018; Ferreira Castro *et al*., 2020; Baltruschat *et al*., 2020). Morphologies, quantified by number of branches, total dendritic length and mean segment length, were similar in synthetic cells grown using the minimum spanning tree (MST) algorithm (Cuntz *et al*., 2010) compared to reconstructions from biological cells (**Figure 2C**). In summary, the algorithm captured the structure of the synthetic trees to such an extent that the morphology could be recovered after removing part of the tree.

A summary of 500 different cuts and synthetic regrowths clearly shows the bimodal distribution between regeneration and invasion (**Figure 2D**). There was a distinct peak at 0% regeneration, *i.e.* 100% invasion, and a flatter distribution of larger percentages of regeneration in the case of the *Drosophila* larval Class IV neurons. There were no obvious relationships between the amount of invasion and model parameters or morphological features. When the results were dissected by the size of the severed branch in *mm* (**Figure 2E**), all types of regrowth were observed for all sizes of severed branches. Only for very large cuts did 100% regeneration become less likely. The exact amount of regeneration depended both on the density of new branches (higher *N*) and on the balancing factor *bf*, the trade-off in the optimal wiring algorithm between minimising the conduction time (*i.e.*, path length) and minimising the total cable length (**Figure 2F**). However, in all cases, both regeneration and invasion were possible outcomes of the synthetic regrowth.

### Repair of different cell types

We then tested whether our regrowth model could be used as a general tool, applicable to a variety of cell types and different species including humans. Our previously established algorithmic generation of distinct dendritic trees of different cell types depends on a single free parameter, the balancing factor (*bf*), weighing material cost (*i.e.* cable length) against conduction time to the soma (*i.e.* path length) (Cuntz *et al*., 2010). Based on recently established algorithms (Bird and Cuntz, 2019), our regrowth model is able to automatically estimate the biological *bf* from any incomplete (input) dendrite morphology. It also analyses the density profile of branch and termination points based on the input neuron to be repaired, and distributes target points accordingly. The MST algorithm (Cuntz *et al*., 2010) then grows new branches along these target points, the number of which is set according to the density of branch and termination points of the input neuron and the size of the growth volume in which the target points are distributed. All parameters can also be adjusted manually. In this way, both highly branched (with low *bf*) as well as less branched morphologies with longer straight branches (with high *bf*) can be modelled.

Examples of such repairs are depicted in **Figure 3**, where panel **A** shows a mouse dentate granule cell (Beining *et al*., 2017). This type of cell minimises predominantly the conduction time (path length) as compared to the material cost (cable length) with a high *bf*. However, the algorithm was also able to repair a mouse Purkinje cell (Chen *et al*., 2013) with many branches. Purkinje cells are known to minimise the material cost more than the conduction time, exhibiting a low *bf*, **Figure 3B**. We also applied the repair algorithm to a spherical synthetic neuron (grown using the MST algorithm of the *TREES toolbox*) that has was cut and then repaired (**Figure 3C**). In this case, we used the conserved growth mode, which limits the regrowth process to the known cut branches. Interestingly, the execution of the procedure from **Figure 2**, where random branches were cut from the morphology and then repaired using the biological regrowth of the *fix tree* function, revealed different distributions of regeneration and invasion in the different cell types (**Figure 3A, B**, *Right*). Although still bimodal, regrowth from the severed branch appeared to be more likely in granule cells when compared to Purkinje cells and *Drosophila* larval Class IV neurons (*c.f.* **Figure 2**). This may be due to the relatively high balancing factor in granule cells.

**Figure 3.**
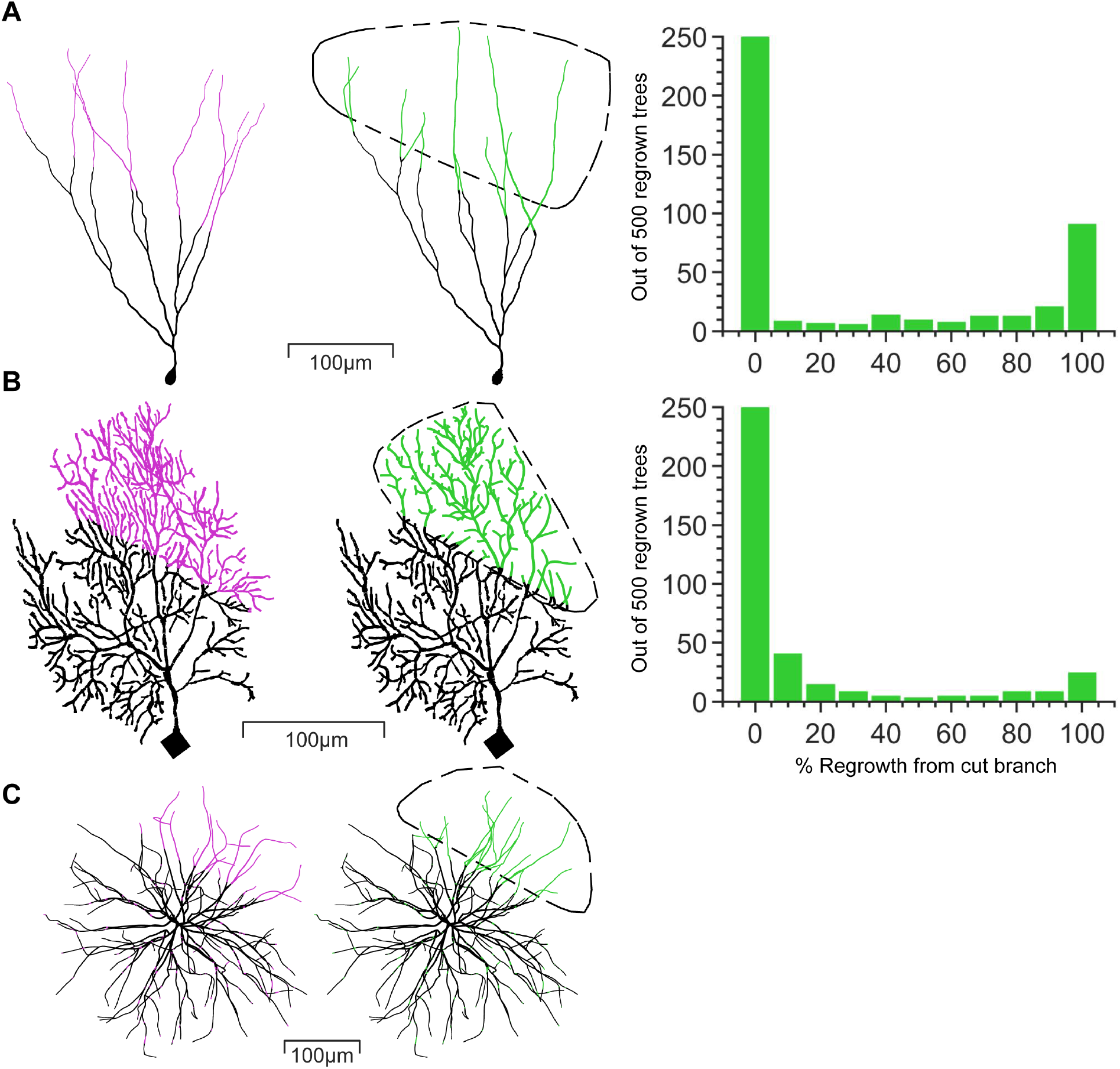
Repair algorithm successfully restores removed dendrites of different cell types with high, low and intermediate balancing factors. **A-C** *Left*, Reference morphology with cut dendrites in magenta, *Middle*, Repaired morphology with restored dendrites in green. The area enclosed by the dashed black line indicates the volume into which the dendrite has grown. **A-B** *Right*, Histogram of 500 regrown morphologies using our repair function `fix tree`, with the percentage of the repair regrowing from the cut branch similar to Figure 2D. **A** Repaired mouse dentate granule cell (Morphology from Beining *et al*., 2017).**B**, Repaired mouse cerebellar Purkinje cell (Morphology from Chen *et al*., 2013). **C**, Repaired synthetic spherical cell created using the `MST tree` algorithm from the *TREES Toolbox* (Cuntz *et al*., 2010).

### Implementation of the regrowth algorithm in a new user interface

Next we used the regrowth algorithm, validated above using the dendrite regeneration data from *Drosophila* da neurons, to develop a new practical tool for repairing lesioned 3D-imaged and reconstructed dendrites. The model was then tested using a dataset of mouse CA1 pyramidal neurons provided by Benavides-Piccione *et al*. (2020) (see more details in supplementary **Figure S1**). The reconstructions of this dataset, like most others (De Schutter and Jaeger, 2000), are incomplete due to the difficult reconstruction process. We have generated an *in silico* model that utilises a graphical user interface (GUI) capable of fixing arbitrary morphologies by adding synthetic dendritic branches to the existing incomplete reconstruction (**Figure 4**). The GUI allows the user to upload any 3D reconstruction and draw or upload any 3D or 2D region where dendrites are missing in the reconstruction. The algorithm then automatically grows the artificial dendrite into the specified volume, preserving the original morphology. This is done by distributing target points in the specified volume and successively connecting them to the input morphology (see Methods). As a reference for the anatomical tissue context, the user can upload a microscope image stack to serve as a background.

**Figure 4.**
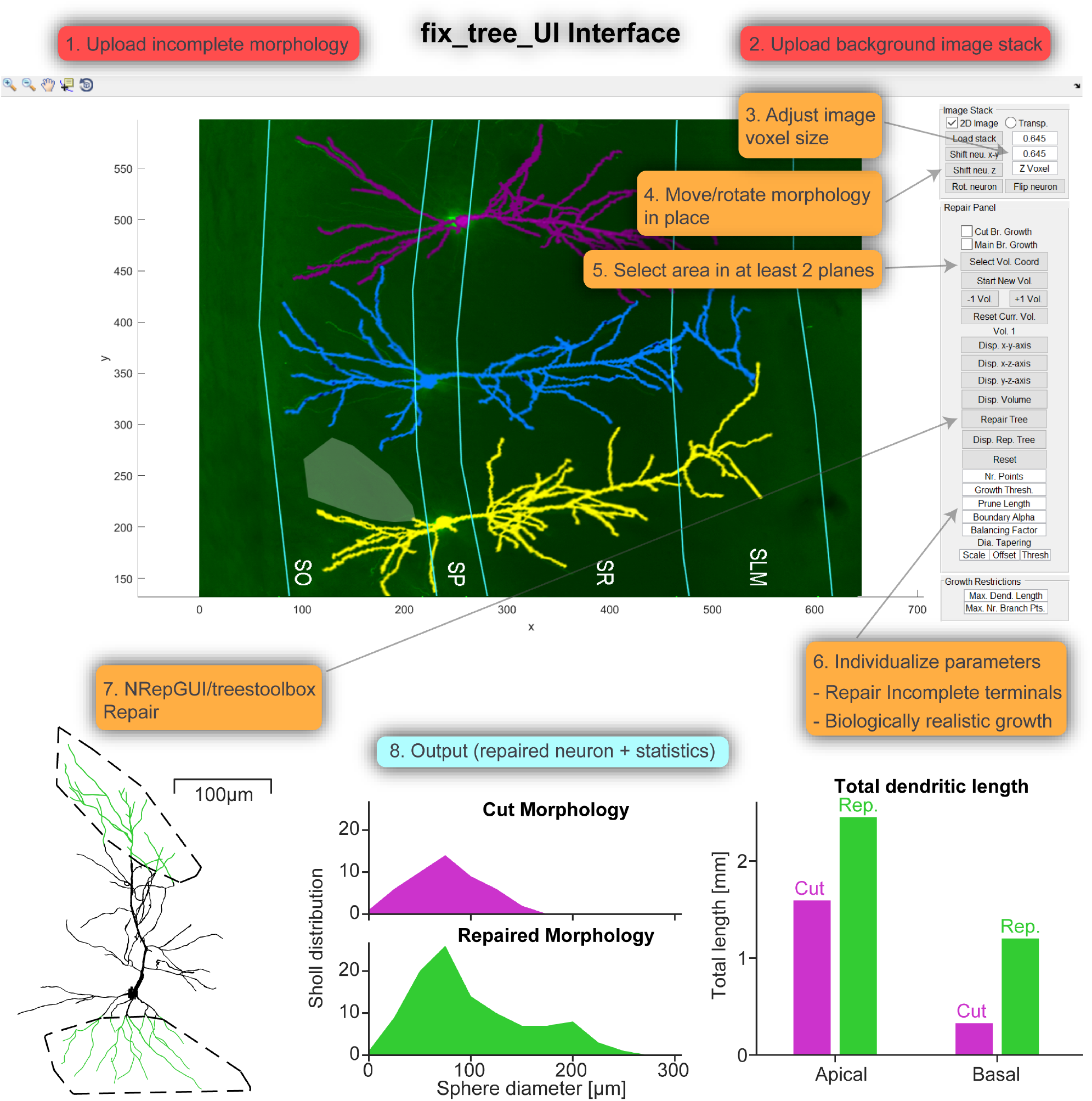
A new software tool for the repair of dendrites with a graphical user interface (GUI) *Top*, Example screenshot of *fix tree UI* (Neuron Repair Graphical-User-Interface). The numbers 1-8 represent the steps of successfully uploading a morphology and background image stack and repairing a missing region. *Bottom*, Showcase of the output of *fix tree UI* with the repaired neuron and two example statistics (the output contains more statistics than shown).

As demonstrated in **Figure 4** *Top*, the image can highlight the different layers of the given brain region, *e.g.* the CA1 region of the hippocampus. This helps as an anatomical indication of where the incomplete morphology might be repaired. The image can be set to the correct size and the morphology moved to the correct location using the image stack panel of the GUI. To draw a 3D target volume, the coordinates for its outline are selected with the cursor in at least two planes (*e.g.* x-y-plane and x-z-plane). Alternatively, the volume coordinates can simply be uploaded. Pressing the Repair button automatically estimates all parameters (see Methods) except the pruning parameters (truncation of terminal dendritic branches below a certain length threshold) and performs the repair. All parameters can also be adjusted manually by the user as well. As shown in **Figure 4** *Bottom*, the GUI outputs the repaired morphology as well as statistical morphological data comparing the input and output morphologies. If available, the user can also upload a reference morphology to be used as a template. The algorithm then matches the statistics of the repair to the reference reconstruction. In this way, the repair mechanism can be tested on sample data before being applied to data from actual incomplete reconstructions.

### Repair of artificially sectioned mouse CA1 pyramidal neurons

Next we tested our repair algorithm on mouse CA1 pyramidal cell morphologies (Benavides-Piccione *et al*., 2020) (**Figure 5**). To assess the quality of our repair algorithm, existing reconstructions were arbitrarily cut at different points and angles in the apical and basal arbour. The original morphology served as a reference and ground truth. The comparison between the reference and the repaired morphology showed the accuracy of the repair (**Figure 5**). Dendritic branching profiles (Sholl, 1953) as a function of the distance from soma showed that the repair algorithm was able to restore the original dendritic shape (**Figure 5**). As the dendrites were intentionally cut, the exact cut-off points were known and the algorithm allowed new dendrites to grow exclusively from these incomplete branches (a forced conserved growth mode).

**Figure 5.**
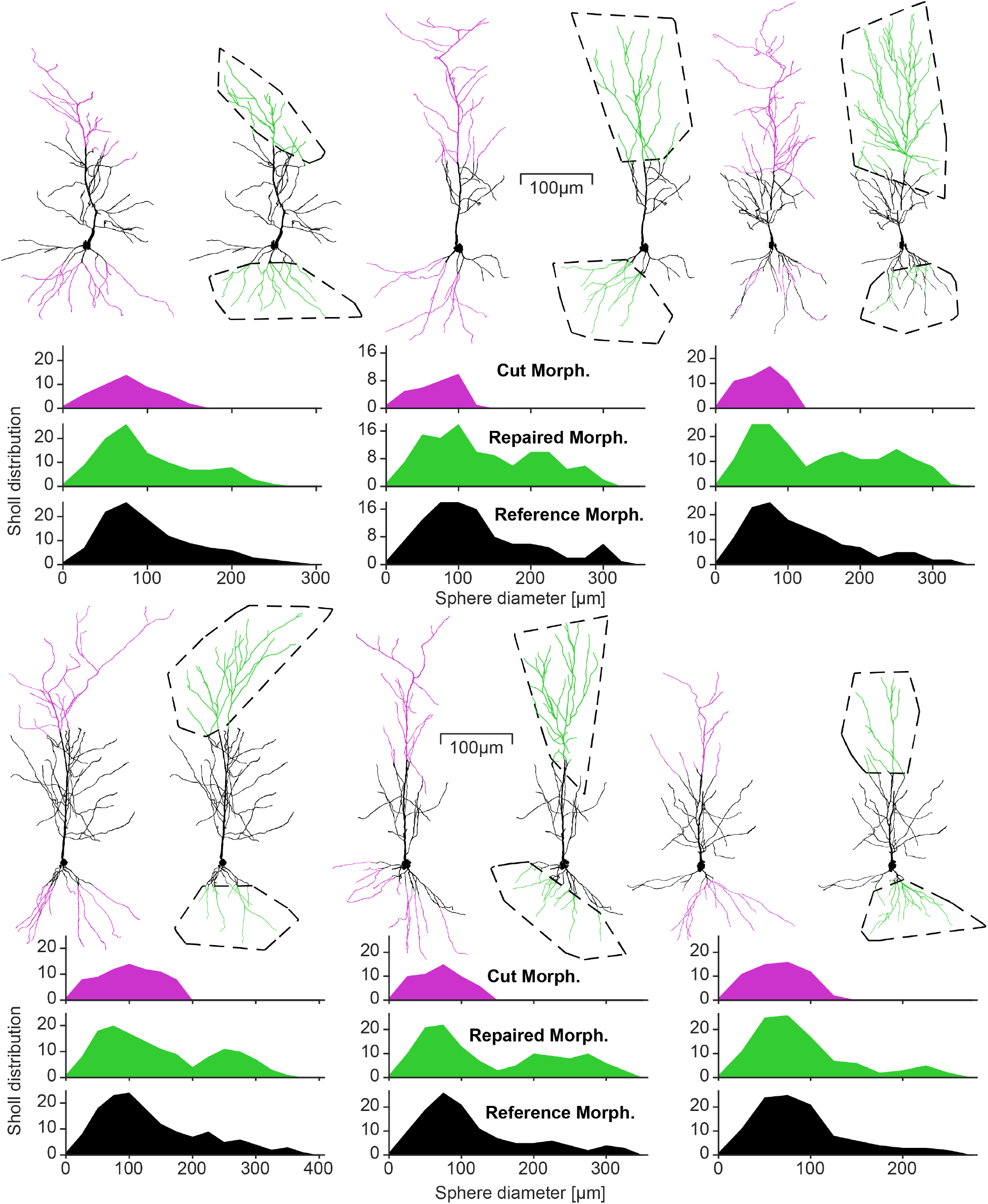
The repair algorithm successfully recovered artificially removed dendrites from mouse CA1 pyramidal cells and restored their Sholl profiles. Six example repairs of apical and basal dendrites of mouse CA1 pyramidal neuron (reconstructions from Benavides-Piccione *et al*., 2020). For each repair, the left morphology is the reconstructed reference with cut branches in magenta and the right morphology is the repaired tree with regrown branches in green. The area enclosed by the dashed black line represents the 3D volume into which the artificial dendrites grew, corresponding to the convex hull of the severed dendrites (see Methods). The graphs below each repair show the distributions of Sholl intersections for the Cut, Repaired and Reference morphologies.

This additional growth mode was inspired by the regeneration observed in biology but was implemented here as a useful option in our software tool. The other mode allows the algorithm to grow new dendrites from any point of the existing morphology, preferably points that are close to the volume chosen for growth (invasion and regeneration). To repair incomplete morphologies we used the conserved growth mode, when the incomplete branches were known, such as in **Figure 5**. This method allows the user to restore a part of the dendrite that they know should be there but could not be reconstructed from their tissue slice. Additionally, since pyramidal cell main apical dendrites can branch, as observed by Benavides-Piccione *et al*. (2020) there is a main growth option for the conserved growth mode (see Methods). With this option, a prominent straight main apical dendrite is grown first and then oblique dendrites are added. As the main apical dendrite is incomplete in all cases shown in **Figure 5**, this option was used for the apical repairs. The extent to which the dendrites grow in a particular direction is given by the growth volume.

From a morphological point of view it is important to accurately analyse the shape and appearance of the neuron as well as the statistics of its morphology. Therefore, **Figure 6** shows further details for the fine-grained morphological statistics of the pyramidal cells from **Figure 5**. The algorithm tries to fit the repairs to exactly match the number of branch points of the reference morphology (**Figure 6**). The model also fits the total dendritic length well in most cases as shown in **Figure 6**, *Top middle*. The remaining four statistics are the dendritic length per segment and the diameter per segment for apical and basal arbours (a segment is measured from one branch point to the next or from a branch point to a termination point). These results show that our model is able to reliably match the morphological properties of mouse CA1 pyramidal neurons in terms of shape and appearance as well as their statistical properties.

**Figure 6.**
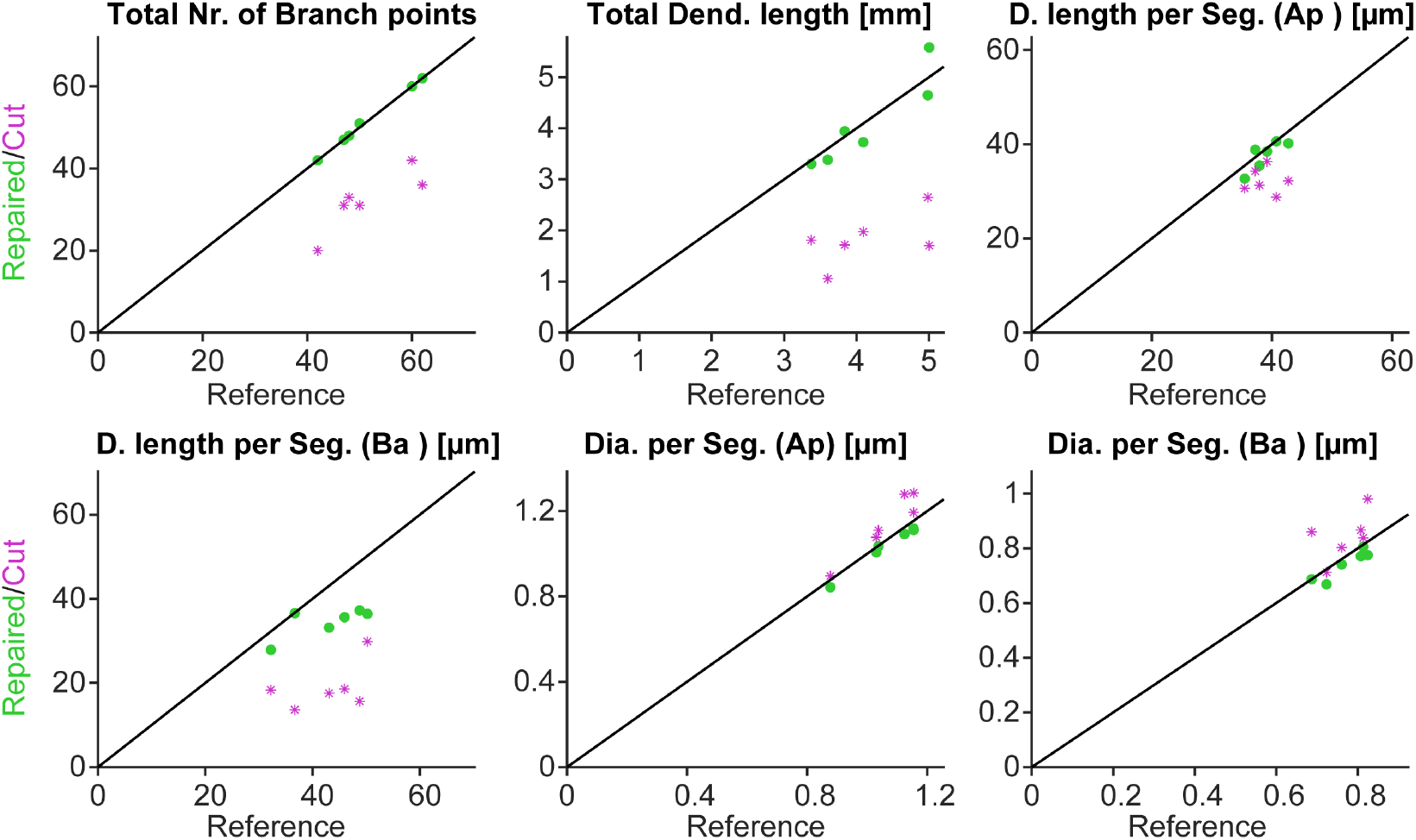
Repair algorithm successfully restores branching statistics of mouse CA1 pyramidal cells. Each graph shows the value of the repaired morphology (green dots) plotted against the value of the original morphology in black on the identity line. For comparison, the data points in magenta show the values for the cut morphologies.

### Repair of human CA1 pyramidal cell reconstructions

We also tested our method on incomplete human CA1 pyramidal neurons. We therefore, applied the repair algorithm to a dataset of CA1 pyramidal neurons from Benavides-Piccione *et al*. (2020) depicted in **Figure 7A,B,C,D**. The morphological reconstructions are shown in panel **B,D**. Similar to the validation process carried out with the mouse reconstructions (**Figure 5**), we then applied our repair algorithm to the original reconstructions from panel **D**. In particular, the basal dendritic arbour and the most distal apical dendritic collaterals and tufts were reconstructed. The results of these extensions are depicted in **Figure 7E**. The dendritic spanning fields of these artificially repaired morphologies are based on the layer limitation boundaries marked out in the slice image. Furthermore, it was assumed that CA1 pyramidal cell dendrites would extend more than halfway into the SLM when the soma of the neuron is close to the SP-SR boundary, in order to make synaptic connections with axons from the perforant pathway (Ito and Schuman, 2012).

**Figure 7.**
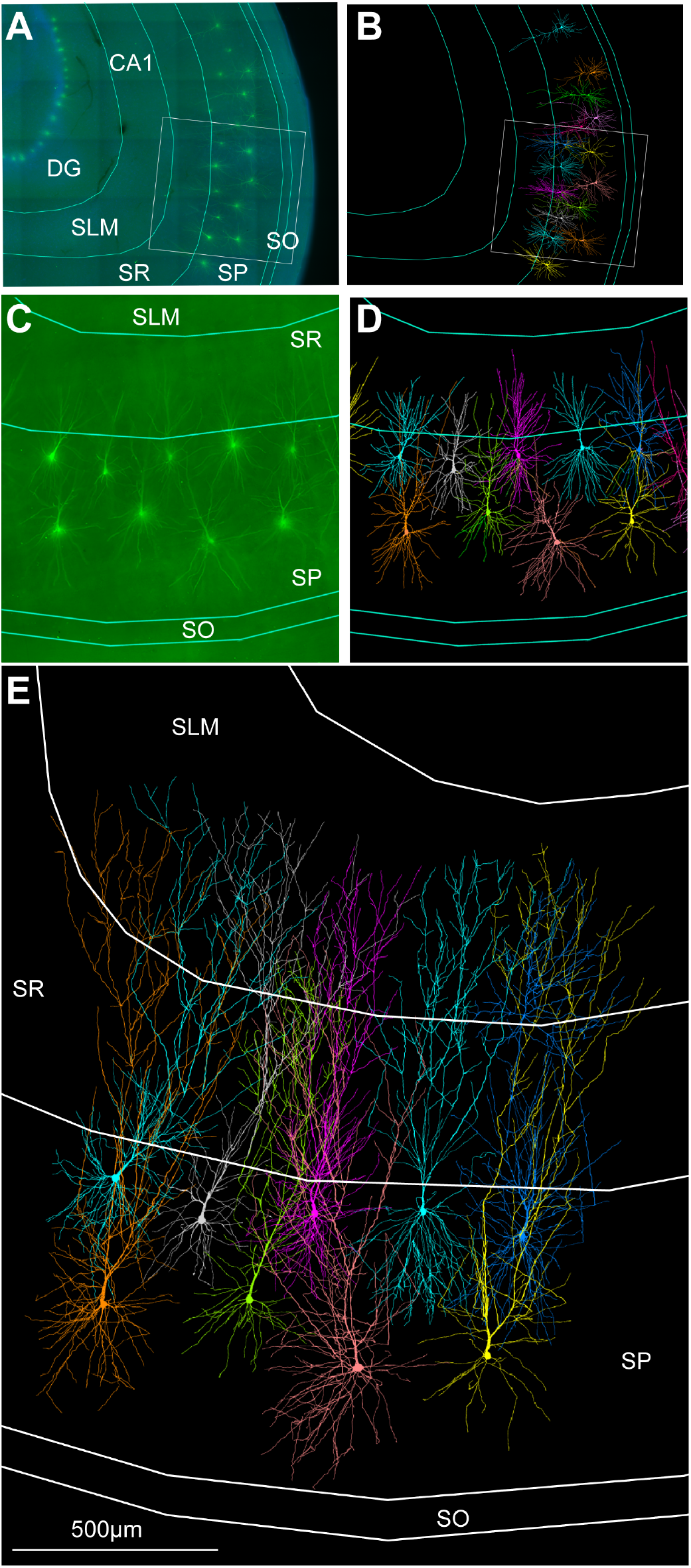
Growth algorithm extends incomplete human CA1 pyramidal cell morphologies. **A**, Confocal microscope image of the human hippocampal CA1 region (DG: dentate gyrus; SLM: stratum lacunosum moleculare; SR: stratum radiatum; SP: stratum pyramidale; SO: stratum oriens) with intracellularly injected pyramidal cells and ROI (region of interest). **B** Morphology reconstructions. **C**, ROI enlarged from **A**. **D**, ROI with overlays of originally reconstructed pyramidal cell morphologies by Benavides-Piccione *et al*. (2020), which are incomplete due to experimental limitations. **E**, ROI showing morphologies from **D** that have been artificially extended in the apical and basal arbour, showing plausible completion of incomplete dendrites based on known layer-specific target growth regions. Each individual neuron has been given a different colour to distinguish the morphologies.

### Restoration of firing behaviour in repaired mouse morphologies and predictions for human data

We tested whether our repair algorithm was able to restore the firing behaviour of the original morphology after regrowth of cut branches (**Figure 8**). We used a biophysical model from Jarsky *et al*. (2005), implemented in mouse (**Figure 8A**) and human (**Figure 8B**) neuron morphologies. Somatic current clamp simulations were performed with the stimulation current increasing in five steps (from 0.16*nA −* 0.24*nA* in **Figure 8A** and 0.26*nA −* 0.46*nA* in **Figure 8B**) and lasting 500*ms* each. The cut neurons clearly displayed hyperexcitable firing behaviour (**Figure 8**). In the repaired neuron, the firing behaviour was restored. We conclude that using our repair tool to restore lost dendritic material can lead to the recovery of the original neuronal excitability, despite incomplete data. **Figure 8C** shows an incomplete human CA1 pyramidal neuron that has been artificially extended (*c.f.* **Figure 7**). The extended version is closer to the actual size of the neuron before the reconstruction process. Consequently, the electrophysiological behaviour predicted for the extended morphology by the Jarsky *et al*. (2005) model differs from the incomplete reference morphology, as excitability is reduced in the extended version (**Figure 8C** *right*).

**Figure 8.**
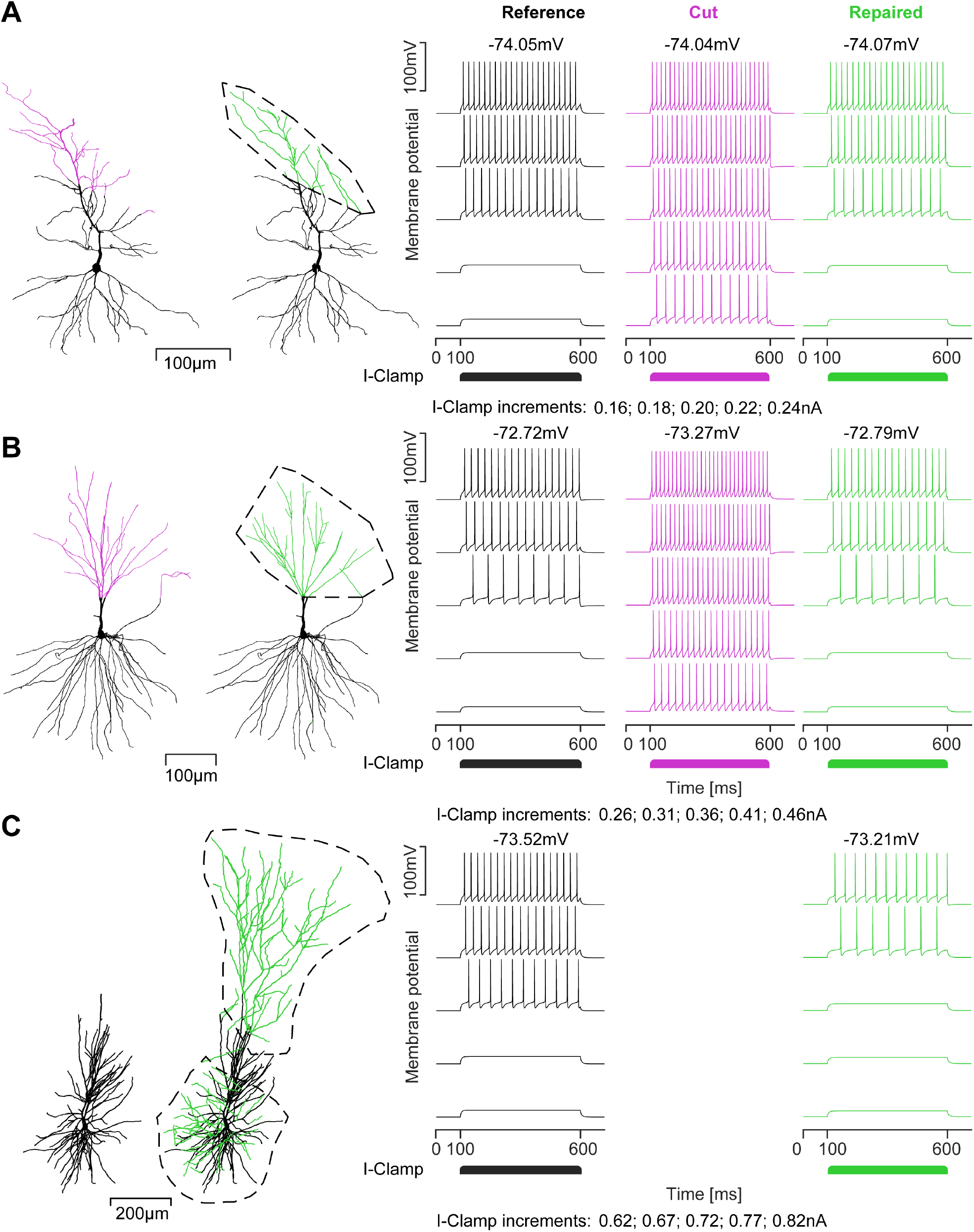
The repair algorithm restores the electrophysiological behaviour of cut and repaired mouse pyramidal cells and allows for better predictions of neuronal function in human neurons. **A**, CA1 pyramidal cell of the mouse. *Left*, Reference morphology, *Middle*, Repaired morphology with growth volume indicated by the black dashed line. Cut dendritic sections in magenta, repaired dendritic sections in green. *Right*, Somatic voltage traces induced by current injections in the soma of reference (black), cut (magenta) and repaired (green) morphology with resting membrane potentials (*Top*) and current clamp increments (*Bottom*). **B**, Human CA1 pyramidal cell. Same arrangement as in **A**. Repair restores the electrophysiological behaviour of the reference neuron. **C**, Prediction of the electrophysiological behaviour of an extended human CA1 pyramidal cell. Arrangement as in **A** but there is no cut neuron since the reference morphology on the left is the full reconstruction as provided by Benavides-Piccione *et al*. (2020), which has been extended (*c.f.* Figure 7).

It has recently been reported that mouse dendrites in cortical pyramidal neurons have lower synaptic thresholds for NMDA spike generation than human dendrites (Testa-Silva *et al*., 2022). To further demonstrate the restorative effects of our repair algorithm on the electrophysiological behaviour of rodent and human dendrites, we performed a computational analysis of their dendritic NMDA spiking. In particular, we were interested in the behaviour of incomplete morphologies that were extended beyond the reconstructed dendritic material (*c.f.* **Figure 7** and **Figure 8C**). In **Figure 9A**, three morphologies were synaptically stimulated in their basal dendrites (highlighted colours; other dendrites in grey) at different Euclidean distances from the soma. The distances were scaled according to the size of the neurons, as the human morphologies were much larger than the mouse morphologies, defined by the percentage of the maximum possible distance within the basal dendrite. Using a passive version of the compartmental model of Jarsky *et al*. (2005) AMPA and NMDA synapses were stimulated. The intensity of the stimulation was determined by the number of synapses distributed over sections of 20*µm*. **Figure 9B** shows example dendritic spike traces with increasing number of synapses, recorded at the site of stimulation, at 85.19% of the maximum possible distance from the soma. We compared a mouse pyramidal cell morphology with an incomplete human reference morphology and an elongated (extended) human morphology. We measured the peak voltage of NMDA spikes evoked by different numbers of synapses at different distances from the soma (**Figure 9C**). For each distance 10 different dendritic locations at that specific distance were tested, as we found a lot of variation in the response (transparent dashed coloured lines) especially close to the soma (**Figure 9C** left). The mouse average peak voltage (solid purple) was generally the largest and had the steepest slope, whereas the voltage peaks in human (solid black) and human extended morphologies (solid green) were similar close to the soma. This is consistent with the findings of Testa-Silva *et al*. (2022), who reported a lower threshold for eliciting NMDA spikes in mouse compared to human layer 2/3 pyramidal neurons. As one moved away from the soma, the response variation decreased in all morphologies with the peak of dendritic spikes in the human reference morphology (black) being more similar to the mouse morphology (purple), whereas the peak of dendritic spikes in the extended human morphology (green) was reduced. Thus, only the repaired human morphology maintained a higher synaptic threshold for NMDA spikes compared to its mouse counterpart. Overall, in agreement with previous findings (Testa-Silva *et al*., 2022), the differences in NMDA spiking were associated with differences in dendritic diameter (**Figure 9D**). Close to the soma, dendritic diameter varied more than further away, resulting in the large variation in NMDA spikes close to the soma. At dendritic locations further from the soma, diameters were consistently larger in the extended human morphology (green) than in the incomplete human morphology (black). With the reduced diameters, the incomplete human morphology (black) showed similar dendritic spikes to the mouse (purple). This indicates that completing a human morphology by extending its dendrites using our repair tool leads to more realistic simulations of NMDA spikes in human neurons.

**Figure 9.**
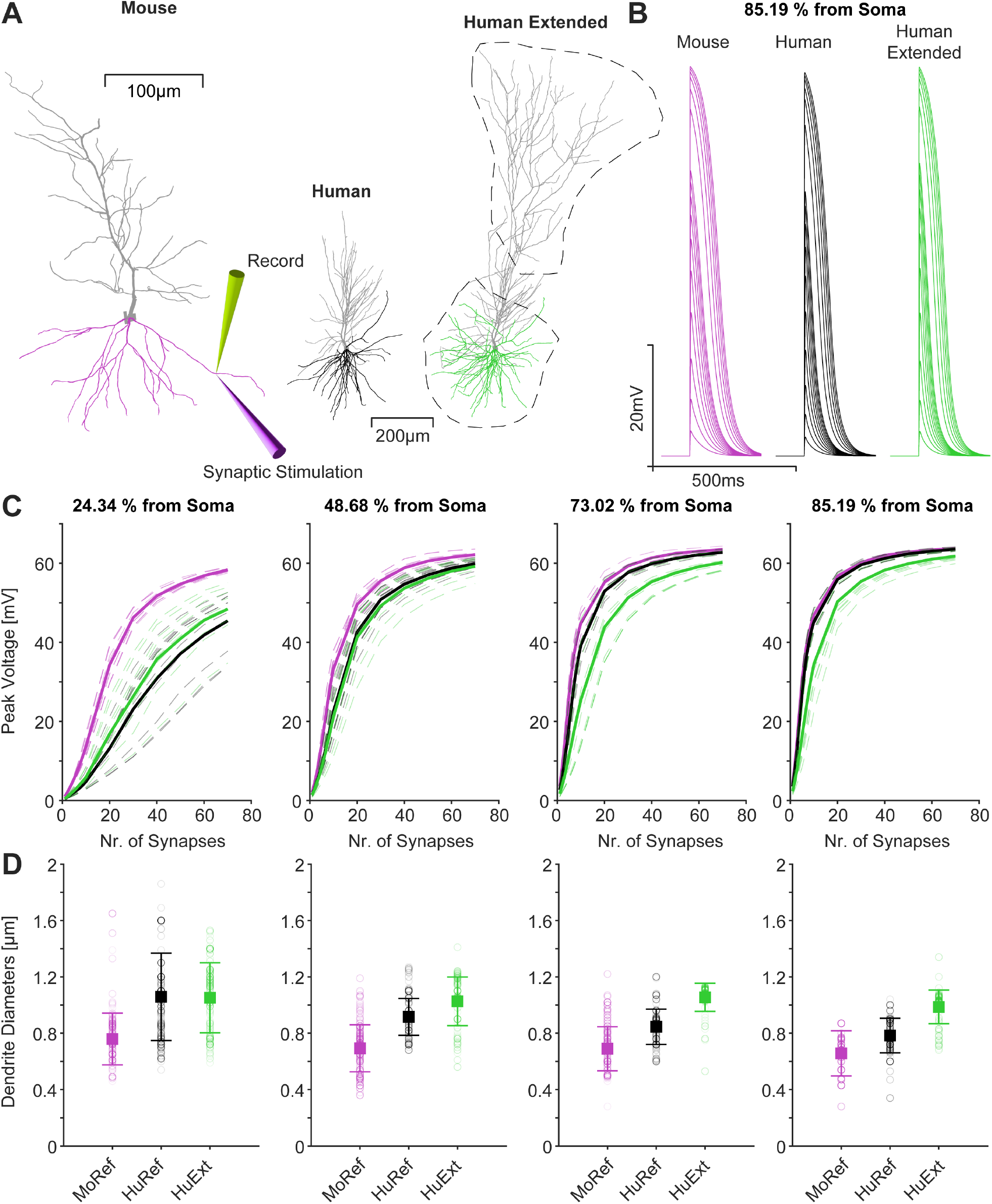
Repairing neuronal dendrites is likely to improve simulations of NMDA spikes, which are reduced in extended human neurons compared to mice. **A**, Mouse CA1 pyramidal cell with basal dendrites in purple. Stimulation and recording sites are indicated on the basal dendrite. *Right*, Human and human extended morphology with basal dendrites in green and black with the growth volumes indicated by the black dashed lines. **B**, Example dendritic NMDA spikes for a mouse (purple), human (black) and human extended (green) morphology at 85.19% of the maximum possible Euclidean distance in the basal tree away from the soma for each morphology respectively. **C**, Peak NMDA spike voltage measured for different numbers of synapses at different distances from the soma in the basal dendrite, given as a percentage of the maximum possible distance in the basal tree (colour scheme as in **B**). For each distance 10 different locations at that distance were tested (transparent dashed coloured lines). The average is shown as a solid line. The synapses were distributed over 20*µm* sections. **D**, Dendritic diameters for the locations described in **C**, with mean and standard deviation.

## Discussion

In this work, we developed a morphological modelling algorithm based on optimal wiring to regrow previously severed dendritic branches. We report four main results. First, we show that the algorithm reproduces an experimentally observed bimodal distribution of dendritic regrowth, consisting of regeneration from lesioned branches and invasion from adjacent branches. Second, when applied to simulated lesions resulting in incomplete 3D morphologies, the repaired dendrites were morphologically similar to the original ones in terms of branching statistics and electrophysiological behaviour. Third, when applied to incompletely reconstructed human CA1 pyramidal neurons, the repair algorithm was able to improve their dendritic structure based on the known anatomical layer-related context. Finally, simulations of species-specific differences in NMDA spiking suggest that our approach will improve predictions of dendritic electrophysiology in incomplete reconstructions.

### Bimodal dendrite regrowth based on the trade-off between optimal cable length and conduction speed

The adapted *TREES toolbox* algorithm (Cuntz *et al*., 2010), which balances material cost, *i.e.* cable length of the dendrite, and conduction time *i.e.* path length to the root (Cuntz *et al*., 2010) was able to successfully regrow dendrites of class IV da neurons of *Drosophila* after removing a part of the tree. The regrown dendrites were statistically similar to the cells under experimental conditions. This indicates that the same balancing factor (which quantifies the trade-off between cable length and conduction speed) underlying the same optimisation algorithm accounts for both a newly generated dendritic tree as well as for the completion of an already existing tree.

Intriguingly, both the computer model and the biological system (Song *et al*., 2012) displayed a binary distribution of invasion vs. regeneration. Song *et al*. (2012) investigated the regenerative capacity of class IV da neurons. Regeneration of class IV dendrites was a commonly observed phenomenon, with 49.4% showing regrowth from the lesioned stem in Song *et al*. (2012) (see also Stone *et al*., 2014). In cells where the severed stem did not regrow, neighbouring branches invaded the area and re-established coverage of the epithelial area by the dendritic network (Song *et al*., 2012). This binary response was clearly seen in both Song *et al*. (2012) and our model. In general, in Stone *et al*. (2014) there was re-coverage of the lesioned area in almost all cases. In the case of 100% invasion, Song *et al*. (2012) reported retraction or stalling of the lesioned dendrite. It was also observed by Stone *et al*. (2014) that if they left a longer stump, the regeneration tended to initiate from there. Therefore it should be investigated how the site of dendriotomy influences invasion versus regeneration. Nevertheless, the close correspondence between the morphological statistics of the original and regenerated nerve cells clearly shows that a dendritic arbour has similar properties before and after the lesion, regardless of whether the empty space is invaded by non-lesioned branches or regenerated from the lesioned stem.

### Human and mammalian dendrite repair

Detailed anatomical data on human neurons remains limited (DeFelipe, 2015). For example, one of the largest public databases of neuronal morphologies, *NeuroMorpho.Org* (Ascoli *et al*., 2007; Parekh and Ascoli, 2013), contains human cell data in only *∼* 4.4% of its entries. Neuroscientists face technical and ethical limitations that limit the acquisition of large datasets from the human brain (Kellmeyer, 2021; Tilimbe, 2019; Palk *et al*., 2020). However, there are structural and functional properties that are specific to the human brain and its neurons (Geschwind and Rakic, 2013; Hofman, 2014; Rilling, 2014; Kaas, 2013; Sherwood *et al*., 2012; DeFelipe, 2011; Oberheim *et al*., 2009; Schmidt and Polleux, 2022), which is why animal neurons cannot completely replace human ones (Zhao and Bhattacharyya, 2018). Human neurons are not only larger but also more complex than those of for example macaques and marmosets (Elston *et al*., 2001). Similar observations have been made when comparing humans and chimpanzees (Bianchi *et al*., 2013). Hodge *et al*. (2019) found a wide range of differences between homologous mouse and human cell types including gene expression, morphology and laminar distribution. To enable more complex brain functions, human neurons have probably evolved special mechanisms such as very strong excitatory synapses, which allow excitatory principal cells to trigger firing in local inhibitory interneurons via a single action potential (Szegedi *et al*., 2016). Recent somatic and dendritic recordings in human neurons and their analyses have also revealed other human-specific electrophysiological properties (Beaulieu-Laroche *et al*., 2021; Moradi Chameh *et al*., 2021; Planert *et al*., 2023; Mihaljević *et al*., 2021; Guet-McCreight *et al*., 2022; Testa-Silva *et al*., 2022; Olah *et al*., 2022; Hunt *et al*., 2023; Szegedi *et al*., 2023; Eyal *et al*., 2018). These species-specific differences may contribute to the unique cognitive abilities of the human brain. Using our approach, such differences could be investigated by first predicting the shape and topology of the putative full morphology reconstruction. In a second step the electrophysiological behaviour and how it differs from the reference reconstruction can be predicted by implementing compartmental models. Extended full human morphology reconstructions could also help to build more accurate human compartmental models. This can be done for any cell type as the *fix tree* function developed in our work is generalised.

Understanding the specific functionality of human neurons requires anatomically complete and reliable datasets of 3D human neuron reconstructions. Our repair tool could help address these issues and alleviate some of the difficulties.

### Restoration of electrophysiological behaviour and practical use for detailed network modelling

As demonstrated in **Figure 8**, cutting off the dendritic arbour of neurons is likely to lead to hyperexcitability in the electrophysiological model, even though the distribution of ion channels is similar in both the cut and the original neuron. The variability in firing behaviour of neurons with similar ion channel distributions has long been recognised (Mainen and Sejnowski, 1996). Neurons that differ only in the geometry of their dendritic arbours produce a wide range of different spiking patterns. Typically, the smaller the neuron, the higher the spiking frequency. Therefore large neurons tend to be less excitable due to their lower input resistance/higher input conductance (Cuntz *et al*., 2021). This is exactly what happens in **Figure 8**, as cutting away some of the dendritic material results in a smaller dendritic arbour, which now has a higher input resistance, thus inducing hyperexcitability.

Compensating for the reduced excitability, large neurons receive more synaptic input (see Cuntz *et al*. (2021)), while small neurons have fewer synapses, reducing the effective current received by the neuron. As we have shown in **Figure 9**, activation of synaptic inputs close to the soma (proximal inputs) leads to large variability in responses, whereas inputs to distal parts of the dendrite appear to produce more consistent dendritic spikes. Therefore, repairing incomplete morphologies and thus restoring distal synaptic input sites may make the electrophysiological behaviour of dendrites more consistent. More importantly, we were able to reproduce the findings of Testa-Silva *et al*. (2022), who showed that the synaptic threshold for NMDA spikes is higher in human pyramidal cells than in mouse pyramidal cells. However, in the case of distal synaptic inputs, the reduced NMDA spike threshold was not present in incomplete human pyramidal cell morphologies. However, when incomplete human dendrites were completed using our repair method (**Figure 9C**), the higher NMDA spike threshold (for distal synapses) was restored. Like Testa-Silva *et al*. (2022), we also found that the differences in the NMDA spike threshold were related to differences in dendritic diameter, which is increased in humans compared to mice. The addition of artificial dendritic material by the repair algorithm increases the average diameter in the distal dendrites of the human extended morphology (green) compared to the incomplete human morphology (black) (*c.f.* **Figure 9D**). This explains the higher threshold for NMDA spike generation in repaired human dendrites than in incomplete human and in mouse dendrites.

In terms of dendritic geometry, not only differences in dendritic length, but also changes in topology such as branching pattern significantly affect the firing behaviour of a neuron (van Elburg and van Ooyen, 2010). Dendritic topology also seems to have an effect on the type of firing, which can be expressed as bursts or regular spike trains (van Ooyen *et al*., 2002). The study by van Elburg and van Ooyen (2010) suggests that changes in the dendritic geometry and topology, which are common in Alzheimer’s disease, epilepsy and mental retardation, have a significant impact on firing behaviour and therefore on information processing and cognitive ability. Therefore, restoring the missing parts of incomplete dendritic morphologies with our repair tool can restore the original firing behaviour of a neuron. The algorithm can be applied to the incomplete morphologies that are available in the databases of the Blue Brain Project (Markram, 2006) and the Allen Brain Atlas Data Portal. Combined with robust and generalisable biophysical models (Beining *et al*., 2017; Cuntz *et al*., 2021), such improved morphologies could be used for large-scale network modelling.

### Limitations of our model and possible extensions

The repair of our software tool is based on the distribution of target points within a selected volume. These points are then successively connected to the existing reconstruction based on wiring optimisation constraints of cable length and conduction speed (Cuntz *et al*., 2010). The volume can be chosen arbitrarily by the user. While this approach is highly flexible and gives the user complete freedom to choose where to grow the morphology, it places an emphasis on the user’s experience, anatomical knowledge and intuition. The possibility to use a suitable reference image helps to assess where the boundaries of an intact morphology are and where certain parts are missing. This facilitates the repair of neurons from brain regions such as CA1 in the hippocampus where the sizes, shapes and anatomical layers are well defined, giving the user a clear indication of where somata are located and where to grow dendrites (as shown in the CA1 region; **Figure 7**). Less well-laminated and defined regions may be more challenging for the context-based neuronal repair. To overcome this problem, data-based predictions for species-, cell-type- and region-specific anatomical boundaries based on morphological statistics would need to be implemented in addition. Such an algorithm would rely on a database containing reconstructions of many morphologies of different cell types, regions and species. To predict the most likely complete boundary of a given input neuron, its cell type and region of origin would have to be specified by the user. Based on the database, an average boundary could be calculated and scaled to the size and dimensions of the input neuron.

As the algorithm uses the uploaded incomplete morphology to automatically determine growth parameters such as the balancing factor, vastly incomplete morphologies can lead to inaccuracies. A morphology with very little dendritic material left is a challenge when trying to estimate growth parameters. Importantly, repaired morphologies can only be used to make predictions. It is important to realise that when you complete a dendritic tree based on the statistics of the remaining tree, you are assuming that the statistics are the same throughout.

### Relationship to other morphological models

While there have been experimental studies investigating how *in vivo* neurons respond to injury and subsequently regrow and repair the damaged dendrites (Song *et al*., 2012; Li *et al*., 2018; Stone *et al*., 2014), artificial repair tools such as Abdellah *et al*. (2018) and *NeuronR* (Coste *et al*., 2021; Anwar *et al*., 2009) mostly focus on removing artefacts that occur during the reconstruction process. Such artefacts include abrupt changes in dendritic thickness at bifurcations, soma profile adjustments, crossing neurites, and dendrites that are disconnected from the soma. Our approach is therefore unique in that it can be generalised and easily applied to any cell type or species easily and is capable of extending the dendritic arbour to create entirely new artificial sections. The easy-to-use graphical user interface allows the repair of incomplete or otherwise unusable morphologies.

Morphological computational models mostly describe the growth as a stochastic process that depends on the branching probability, the number of branching events and the number of segments (van Pelt and Schierwagen, 2004; Ascoli and Krichmar, 2000; Donohue and Ascoli, 2008). It has recently been shown that a sequential stochastic growth and retraction algorithm is able to generate dendritic trees of *Drosophila* larval sensory neurons that are realistic in terms of both function and optimal wiring (Baltruschat *et al*., 2020; Ferreira Castro *et al*., 2020), see also Palavalli *et al*. (2021). Similarly, building on the *TREES toolbox* (Cuntz *et al*., 2011), our repair tool also takes wiring optimisation into account. Therefore switching between different cell types with different wiring constraints can be done by adjusting a single free parameter, the balancing factor *bf*, which determines the cell type specific optimal balance between cable length and conduction speed. Using a limited set of parameters is the best way to implement a model if one wants to avoid overfitting problems (Poirazi and Papoutsi, 2020). This simplicity makes our tool adaptable and easy to generalise to different morphologies and helps to understand whether certain cell types optimise their dendrites primarily for material or conduction costs.

## Conclusion

The *TREES toolbox*, extended by the new `fix tree` function, allows for a range of investigations of dendritic anatomy, both during growth and repair, using synthetically grown dendritic structures. The morphological, and by extension functional, changes following cut and repair have not been extensively studied *in vivo*, and can be addressed *in silico* using our repair tool for both synthetic cell models and biological reconstructions. By making this tool widely available to the scientific community, datasets of human neuronal reconstructions could be improved and expanded. Such datasets could eventually provide the insight we need to understand what makes the human brain different from other species.

## Materials and methods

### Regrowth of lesioned class IV da-neurons of *Drosophila melanogaster*

We reconstructed the lesion paradigm, regrew the missing branches to re-cover the target area of the cell, and assessed the differences in morphology using statistical parameters. To study the bimodal distribution of regeneration from the lesioned stem and invasion we severed random dendritic subtrees of *Drosophila* da neurons, Purkinje cells and granule cells with lengths between 50*µm < L <* 1, 000*µm* for 500 trials. Using the repair tool we regrew these 500 morphologies based on the volume previously occupied by the cut branches. To avoid a bias toward regeneration or invasion, target points were distributed within the growth volume with a given margin of *R_d_* away from any point of the lesioned neuron. To assess the distribution of regrowth, we determined what percentage of the regrown dendritic material was regenerated from the lesioned stem. The different growth modes of the GUI, and in particular the `fix tree` function that is at the heart of the repair tool, are described in more detail in the next section.

### The fix tree function of the repair algorithm

Based on the regrowth algorithm for *Drosophila* neurons (see above), we developed a stochastic model of regrowth after dendritic lesions in mouse and human CA1 pyramidal neurons using custom code implemented in the MATLAB-based *TREES toolbox* (Cuntz *et al*., 2011).

The repair algorithm is based on the minimum spanning tree (MST) function (`MST tree`) from the *TREES toolbox* (Cuntz *et al*., 2010). A tree is the representation of the morphology of a neuron by a set of nodes and an adjacency matrix defining the connections between these nodes. The distance between two consecutive nodes was adjusted by resampling the tree to achieve a distance between neighbouring nodes of 1*µm* without significantly changing the branching morphology. The missing dendrites were regrown by distributing the target points over an area/volume *V*, which is an input to the function. To match the clustering of branch and termination points in the input neuron, the density profile of its spanning field is analysed and random clustered points are distributed accordingly using a Monte Carlo approach (available in the *TREES toolbox*). The number of target points *Npts* required is estimated by evaluating the density of branch points in the input neuron along with the size of the area/volume *V*. `MST tree` then connects these points successively to the existing input neuron using a cost function (see Cuntz *et al*., 2010) that depends on the balancing factor *bf*, which weights the conduction time (path length cost) against the material cost (wiring cost).

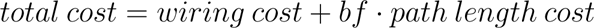

The balancing factor *bf* is estimated by analysing the original input morphology using the *bf tree* function in the *TREES toolbox* (Bird and Cuntz, 2019). The maximum distance a single connection can span is limited by the growth threshold *G_thr_*, which is calculated by measuring the part of a straight line *m*, passing through the neuron root *R* (soma) and the point lying between the mean volume coordinate *V_mean_* and the volume coordinate furthest away from the root node *V_far_*, that lies within *V*.

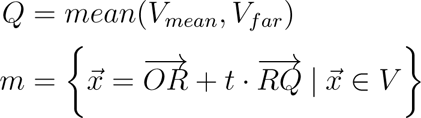

The values of *t* must be chosen so that *m* lies within *V*. New dendrites can grow from any point in the input tree within the range of *G_thr_* (biological growth). This is the first of two growth modes available in the `fix tree` function, which fills the space by growing from lesioned or intact parts of the dendritic arbour. Alternatively, the algorithm can grow new dendrites exclusively from incomplete terminals of the neuron’s branches (conserved growth), repairing a missing part of a severed neuron. Such incomplete terminals must be specified by the user with their exact coordinates in the uploaded morphology file. The algorithm only selects incomplete terminals for growth that are in close proximity to the growth area/volume *V*. The maximum distance an incomplete end can have to *V* depends on the size of the original input tree. Additionally, the noninvasive/conserved growth mode has an option (main growth) specifically designed for severed apical dendrites of pyramidal neurons, since they usually feature one or more prominent main apical dendrites (Benavides-Piccione *et al*., 2020). These grow approximately in a straight line from the root of the tree. If this option is enabled, the algorithm will determine the thickest incomplete terminals in relation to all incomplete terminals and grow a main branch from these first, up to approximately 95% of the length of the growth volume. The direction and distance the main branch will grow is estimated by the same straight line *m* that was calculated earlier. *m* serves as a template for the main apical branch. The algorithm then proceeds as before, allowing dendrites to branch from the newly added main apical section.

In addition to the input neuron to be repaired, a reference morphology (if available) can be passed to the function. The algorithm then matches the number of branch points *NBr* of the repaired neuron to *NBr* in the reference neuron or to an arbitrary number (greater than *NBr* in the input neuron) by iterating over the growth process but successively adding more target points until the desired number is reached.

The area/volume *V* for the repair dendrites to grow into is an input to the function and can be any set of user-defined 2D or 3D points. The volume is then defined by using the boundary function in *MATLAB*, which uses *α*-shapes (Akkiraju *et al*., 1995) to determine the outline of a set of points. How tightly the boundary fits is determined by a single parameter *α*, where *α* = 0 is the convex hull and *α* = 1 is the tightest boundary.

To better match the appearance of the existing input neuron, low-pass filtered spatial noise is imposed on the coordinates of the grown dendrite as a spatial jitter. To achieve realistic diameter values for the grown dendrites, a quadratic taper is applied using the quadratic tapering algorithm of the *TREES toolbox* developed by(Bird and Cuntz, 2016). The taper parameters are estimated based on the original existing morphology reconstructions. The repaired morphology is then tapered using these estimated parameters scaling down towards a minimum diameter in the terminal branches of the morphology as proposed by Liao *et al*. (2021). Since towards the very tips of the dendrites the diameters level off to a constant value, depending on the species, any diameters that fall below an adjustable threshold are set to that threshold value. Optionally, the morphology can be pruned to a desired dendritic length (*e.g.* length of a reference morphology) by first matching *NBr* and then trimming any excess material. By default, all parameters are estimated by analysing the morphology of the input neuron. The main parameter of the `MST tree` function, the balancing factor *bf*, is estimated by analysing the root angle distribution as introduced by Bird and Cuntz (2019).

The GUI fix tree UI, for easy access to the fix tree function, was programmed in the GUIDE MATLAB environment with a custom design interface (see **Figure 4**).

### Electrophysiology (*T2N*)

For electrophysiological compartmental modelling we used the previously developed *T2N* (*TREES*-to-*NEURON*) software interface (Beining *et al*., 2017) in MATLAB which links the compartmental modelling package *NEURON* (Carnevale and Hines, 2006) and the *TREES toolbox*. *T2N* allows for the creation and use of existing complex electrophysiology models, many of which are readily available from https://senselab.med.yale.edu/modeldb (McDougal *et al*., 2017). Any morphology in the *TREES Toolbox* can be uploaded to *T2N* and is then equipped with ion channel conductances specified by the biophysical model. We simulated somatic current injections with a duration of 500*ms* and ramping intensity for both mouse and human morphologies. Current clamps were performed on the reference, the repaired and the artificially cut morphologies respectively in order to compare their behaviour. We used a biophysical model from Jarsky *et al*. (2005), previously imported into *T2N*. The model by Jarsky *et al*. (2005) incorporates four active voltage channels (conductances). These channels include the following: a voltage-gated Na^+^ channel, a delayed rectifier K^+^ channel, a distal A-type K^+^ channel with an elevated half-inactivation voltage and a proximal A-type K^+^ channel. The model distributes these ion channels along the dendrites as a function of the length of the direct path to the soma. The model of Jarsky *et al*. (2005) includes a weak excitability version that follows a uniform distribution, which was used to model the delayed rectifier K^+^ and the Na^+^ channel. Following the experimentally reported sixfold increase in conductance along the apical dendrites, the A-type K^+^ current was modelled accordingly. The result is linearly increasing slopes of variable nature between soma and tuft for different morphologies. The regions of the apical dendrites were defined as follows: the boundaries for the apical trunk (proximal apical) were set to contain 3.14% of the total apical length. The medial apical dendrites contain 36.27%, the distal 68.90% and the tuft 100% of the total apical length. The dendrites were divided at path distances of approximately 100*µm*, 300*µm* and 500*µm*. To simulate synaptic dendritic spikes, we implemented AMPA and NMDA synapses at different locations on the basal dendrites of three morphologies (mouse, human and human extended). The simulations were again carried out using the model of Jarsky *et al*. (2005), but with all active ion channel conductances switched off, leaving only the passive properties of the model. The dendritic diameters on these morphologies were adjusted to eliminate any artefacts that arise during the reconstruction process when using Neurolucida 360 (MBF Bioscience). Synaptic stimulation was carried out at different euclidean distances from the soma based on the maximum possible euclidean distance from the soma of the basal dendrite. The distances were thus scaled for the different morphologies respectively, since the human morphologies are much larger than mouse morphologies. The procedure is designed to expand on what was previously done by Testa-Silva *et al*. (2022), who measured NMDA spikes in human and mouse layer 2/3 pyramidal neurons at only one fixed distance (150*µm* from the soma) for mouse and human. To account for morphological variability, 10 different sites were simulated for each distance and the average was calculated. The stimulation strength was determined by the number of synapses, which were distributed over segments of 20*µm* length. We then recorded the dendritic spike response for different numbers of synapses with *gAMPA* = 25*pS* and *gNMDA* = 500*pS* at the stimulation site. We also measured the diameters for each part of the 20*µm* sections.

## Acknowledgements

We are grateful to Y. Song for communications in the initial phase of the project and to Martin Mittag for his support in NMDA spike simulations. This work was supported by BMBF grants (No. 031L0229 – HUMANEUROMOD to P.J.; No. 01GQ1406 — Bernstein Award 2013 to H.C.), a DFG grant (CU 217/2-1) and the von Behring Rö ntgen Foundation (to P.J.). The authors declare to have no competing financial interests.

## Author contributions

M.G., H.M.M., B.S., J.D., R.B.-P., H.C., P.J. designed the study. M.G. performed the simulations and analysed the data. M.G., H.M.M., B.S., J.D., R.B.-P., H.C., P.J. wrote the paper.

## Supporting information

**Figure S1.**
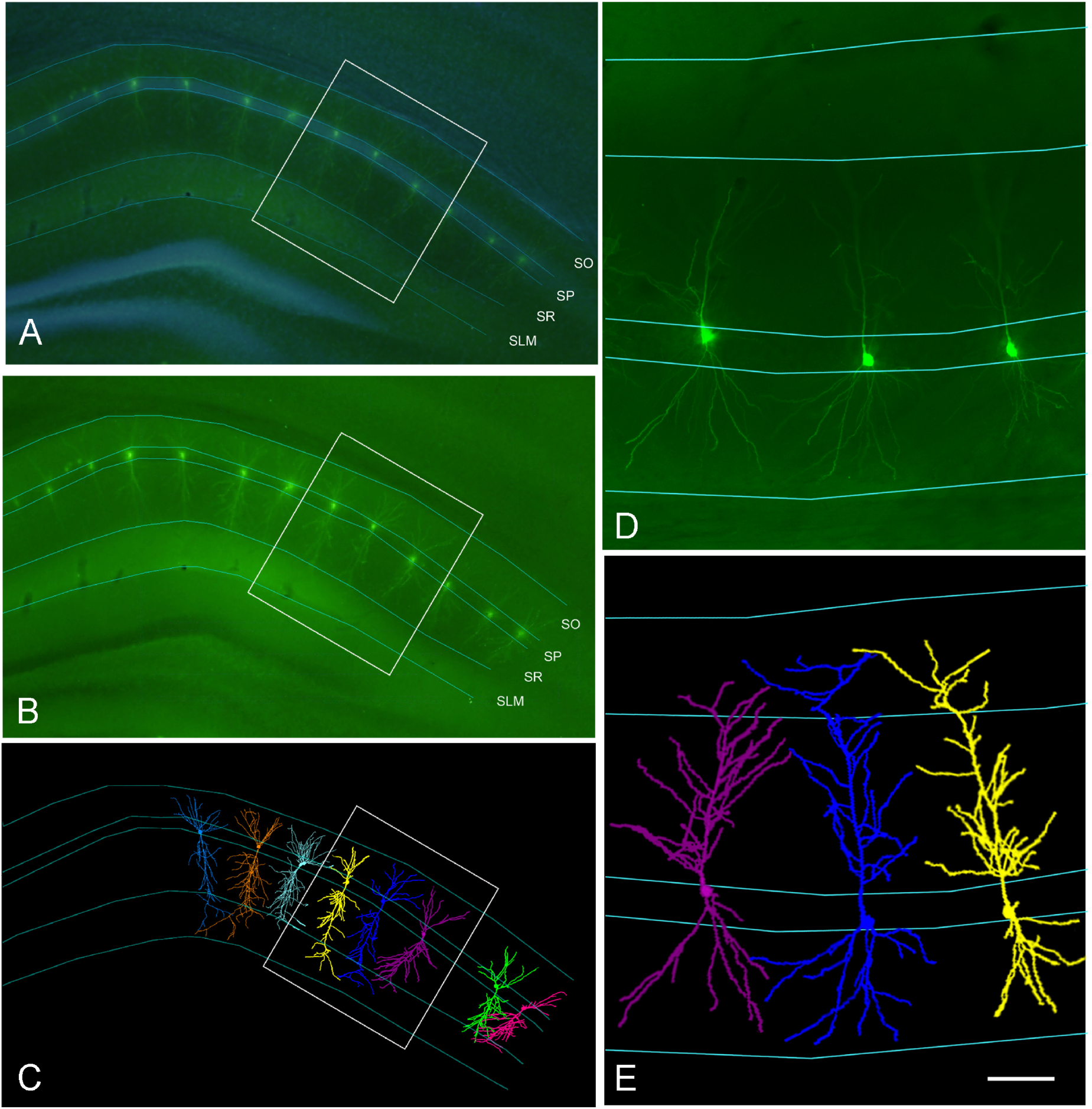
Hippocampal CA1 region of the mouse. **A,B**, Confocal microscope image of the mouse hippocampus with stained pyramidal neuron morphologies, region of interest (ROI) and marked layers (SLM: stratum lacunosum moleculare, SR: stratum radiatum, SP: stratum pyramidale, SO: stratum oriens). **C**, Morphology reconstruction overlays with marked layers. **D**, Magnified ROI with marked layers. *E*, Magnified ROI with marked layers and example of reconstructed morphology overlay. Imaging data were taken from Benavides-Piccione *et al*. (2020).

